# Anionic bacterial sphingolipids increase membrane stiffness

**DOI:** 10.64898/2026.06.07.730480

**Authors:** Joshua D. Chamberlain, Jesse W. Sandberg, Ziqiang Guan, Benjamin P. Bratton, Grace Brannigan, Eric A. Klein

## Abstract

Recent genetic and bioinformatic studies have led to the discovery that many bacterial species encode the genes required to produce sphingolipids. Shotgun lipidomic studies have identified numerous sphingolipid species with novel structures that do not exist in eukaryotic organisms. The impacts of these lipids on the biophysical properties of bacterial membranes have not yet been determined. In this study, we purify a novel anionic bacterial sphingolipid, ceramide phosphoglycerate (CPG), and investigate its effect on membrane zeta potential and bending stiffness. CPG and its precursor, ceramide 1-phosphate (C1P), are shown to increase the magnitude of the membrane zeta potential. These sphingolipids also increase the stiffness of these membranes, with CPG increasing rigidity more than C1P or ceramide. This work provides experimental and computational methods of lipid isolation and characterization that may be broadly applicable to a variety of uncharacterized bacterial sphingolipids.

**SIGNIFICANCE:** The diversity of bacterial sphingolipids far exceeds those found in eukaryotes. However, the function and biophysical properties of these lipids are unknown. Characterization of these lipids is a challenge as they are not commercially available. In this study, we developed experimental methods to purify the anionic sphingolipid ceramide phosphoglycerate and incorporate it into liposomes for analysis. Furthermore, we built computational tools to determine the bending stiffness of sphingolipid-containing vesicles from thermal fluctuation data.

## INTRODUCTION

The membrane bending modulus *k*_*C*_ is a fundamental material property underlying the deformability and mechanical behavior of membrane-bound systems. In cell biology, the bending stiffness constrains diverse membrane remodeling processes including budding, fission, fusion, trafficking, and cellular motility (1–5) (and citations therein). In engineered lipid nanoparticles (LNPs), membrane stiffness modulates LNP-membrane fusion, endosomal escape, and cellular uptake (6–11). As bending stiffness is directly related to the molecular composition of the membrane, establishing quantitative relationships between lipid molecular features and membrane bending rigidity remains an important challenge.

Initial characterization of membrane stiffness has largely focused on phospholipid bilayers, where several factors have emerged as critical regulators of bending rigidity, primarily the membrane thickness and lateral packing and chain order of the lipids. These factors are most often mediated by acyl chain length and saturation (4, 12). Lipids with large polar headgroups can increase the surface charge density, creating repulsion between headgroups and inducing curvature in the membrane, affecting the membrane’s compressibility modulus and area per lipid (13). More recently, membrane characterization has expanded to study the impact of sphingolipids on bending stiffness. Sphingomyelin-containing membranes become more flexible with the addition of cholesterol (14, 15), and the enzymatic conversion of sphingomyelin to ceramide within phospholipid membranes causes phase-separation, increasing the bending rigidity in domains containing ceramide (16). To date, the vast majority of studies on membrane bending have focused on those containing eukaryotic lipid species.

By contrast, the viscoelastic properties of membranes containing bacterial lipids are largely understudied. In particular, bacteria synthesize an extraordinary array of uncharacterized lipids, often in a species-specific manner, many of which have unknown physiological roles. The production of these lipids is often regulated by environmental factors, suggesting that they may help mediate cellular adaptation. We recently demonstrated that *Caulobacter crescentus* produces the novel lipid ceramide phosphoglycerate (CPG), which enables bacterial survival in the absence of the otherwise essential outer-membrane component lipopolysaccharide (17).

Given the physiological impact of CPG and its negatively-charged headgroup, we chose this lipid as a platform for developing methods for bacterial lipid characterization. As shown in Fig. 1A, bacterially-derived CPG contains several molecular features suggesting that it might stiffen membranes: a highly charged headgroup, a sphingosine backbone, a mono-unsaturated acyl chain, and acyl chain hydroxylation. Given the lipid charge and expected stiffness, new experimental and computational tools were required for lipid purification and bending stiffness determination, respectively. In this work, we assessed the impact of CPG on membrane bending modulus and zeta potential. We found that CPG can increase bending stiffness 4-fold over neutral ceramide and increasing CPG content decreases zeta potential monotonically.

**Figure 1:**
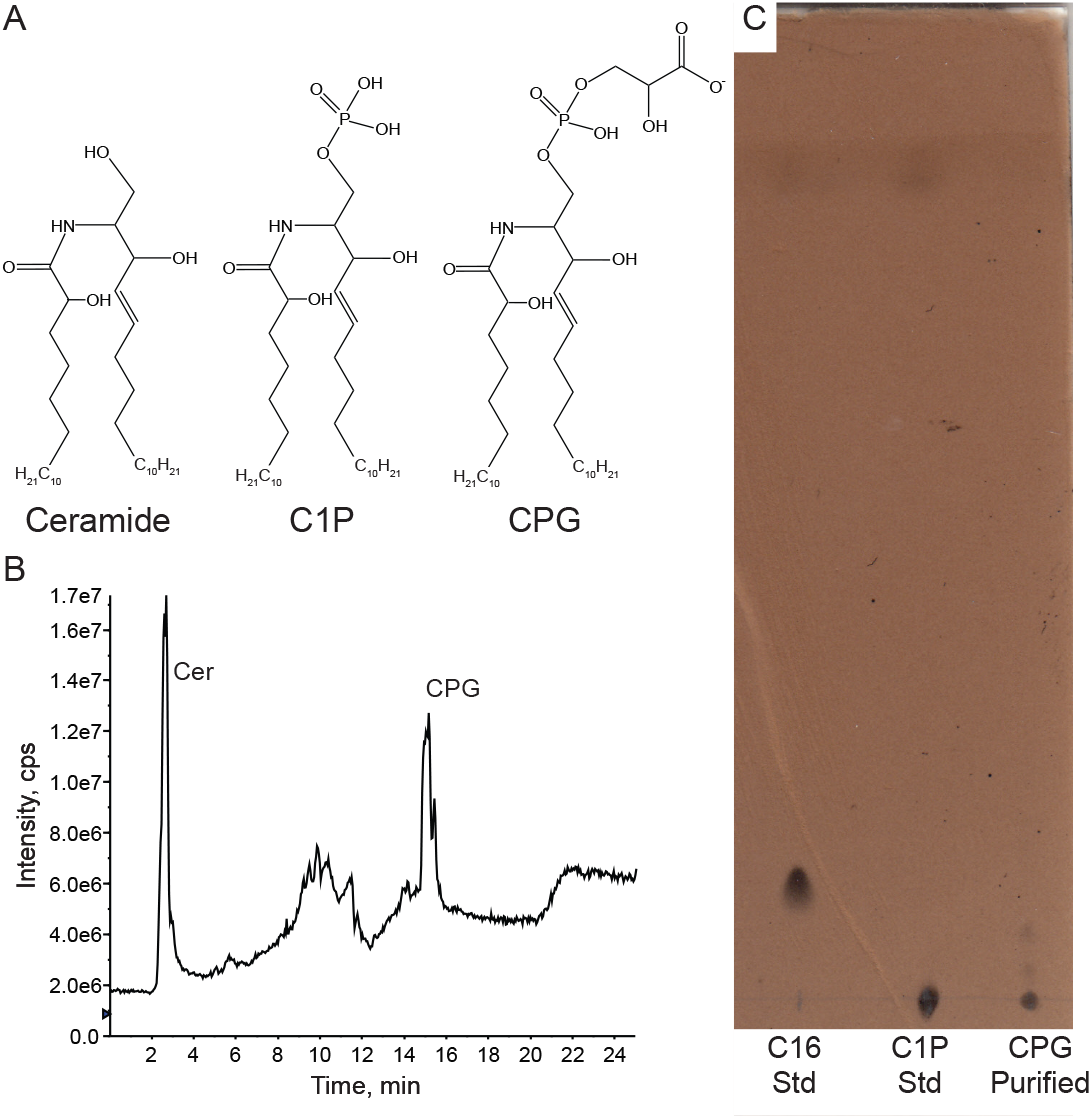
Lipid extraction and purification process to isolate anionic bacterial CPG. (A) Molecular structures of ceramide, C1P, and CPG. (B) Total ion chromatogram of the purified CPG lipid. (C) TLC comparing commercial ceramide (Cer) and C1P with extracted CPG. When run in tandem on a TLC in solvent B and charred with sulfuric acid spray, ceramide (lane 1) runs higher than C1P (lane 2) or CPG (lane 3). The lack of a spot in the CPG lane with the same migration distance as ceramide suggests that ceramide peak observed in (B) is due to decomposition of the CPG headgroup during LC/MS analysis.

## MATERIALS AND METHODS

### Lipids

All commercially available lipids were acquired from Cayman Chemicals (Ann Arbor, MI): C16 ceramide (d18:1/16:0, 10681); 1,2-dioleoyl-sn-glycero-3-phosphatidylcholine (DOPC, 15098); 1-palmitoyl-2-oleoyl-sn-glycero-3-phosphatidylcholine (POPC, 15102); C16 ceramide-1-phosphate (d18:1/16:0, ammonium salt, 22542). Ceramide phosphoglycerate (CPG) was extracted and purified from *Caulobacter crescentus* as described below.

### TLC solvents

Solvent A was CHCl_3_:MeOH:acetic acid:H_2_O (8:3:2:1, v/v) and was used for preparative TLCs in the purification of CPG. Solvent B was CHCl_3_:MeOH:acetic acid (190:9:1, v/v) and was used for analytical TLCs comparing commercial vs purified sphingolipids.

### Lipid extraction

Ceramide phosphoglycerate (CPG) was extracted, enriched, and purified from a *C. crescentus* strain with the genotype Δ*ccna_01217, PvanA ::His-ccna_01219*. The deletion of *ccna_01217* prevents the conversion of CPG to CPG2 (17). Overexpression of *ccna_01219* enhances production of CPG. Cultures of this strain were grown in 20 liters of peptone yeast-extract medium (PYE) containing 2 g/L peptone, 1 g/L yeast extract, 1 mM MgSO_4_, and 0.5 mM CaCl_2_. Induction of *ccna_01219* expression was initiated by adding 0.5 mM vanillate. Initial attempts to extract CPG using the traditional method of Bligh and Dyer (18) resulted in a poor yield, likely due to the polar nature of the lipid headgroup (Figure S1A). An alternative approach to maximize the yield of CPG was developed. Briefly, 20 liters of cells were pelleted and resuspended in 150 mL of molecular biology grade water. Concentrated cells were lysed via French press at 20,000 PSI and centrifuged at 67,000× *g* for 1 hr to isolate the bacterial membranes. Membranes were then resuspended in 50 mL molecular biology grade water before vigorously mixing 1 volume of sample with 3.75 volumes of 1:2 (v/v) chloroform (CHCl_3_):methanol (MeOH). 1.25 volumes each of CHCl_3_ and water were added sequentially with vortexing. The extraction mixture was then acidified below the pKa of CPG to a pH of 2 with hydrochloric acid (HCl) before vortexing, centrifugation at 7,000× *g* for 5 minutes, and isolating the lower organic phase with a glass Pasteur pipette, resulting in a sample enriched with CPG (Figure S1B). Upon completing the extraction and evaporating the remaining solvent with argon, phospholipids in the sample were hydrolyzed by resuspending the lipids in MeOH, 0.1 M potassium hydroxide (KOH), for 4 hours before neutralizing and re-extracting via acidic Bligh-Dyer as previously described (Fig S1C). Lipid samples were applied to TLC plates (Sigma-Aldrich, 10 ×20 cm, TLC silica gel 60, Product 1056260001) and developed in solvent A. TLC plates were then dried and lipids were visualized by exposure to iodine vapor in a sealed chamber. The lipids of interest were scraped off with a clean razor and recovered from the silica via an additional acidic Bligh-Dyer extraction as described. The composition of the purified sample was validated by normal phase liquid chromatography/mass spectrometry (LC/MS) in the negative ion mode as previously described (Figure 1B) (19, 20). We noted the presence of a peak corresponding to ceramide; however, this is likely a decomposition product of CPG, as indicated by the lack of a corresponding TLC signal when compared with a ceramide standard (Fig 1C).

### Analytical TLC

Lipid samples were applied to TLC plates and developed by running twice in Solvent B. TLC plates were then dried and evenly sprayed with 4.2 M H_2_SO_4_, dried, and charred on a hot plate. The slides were kept at room temperature for 24 hours prior to imaging to improve the sample contrast.

### SUV formation

Small unilamellar vesicles (SUVs) were formed via solvent injection. Lipid formulations were solubilized at 1 mg/mL in tetrahydrofuran (THF) and injected dropwise into stirred dispersant buffer (either Dulbecco’s Phosphate Buffered Saline (DPBS) or 10 mM HEPES, pH 7). THF was removed by warming the solution above the boiling point of THF (67 °C) resulting in a final lipid concentration of 125 *μ*g/mL. Size and polydispersity were measured via dynamic light scattering (DLS; Malvern Zetasizer Nano-ZS).

### GUV formation

Giant unilamellar vesicles (GUVs) were formed via electroformation. Lipid formulations were solubilized in CHCl_3_ to 5 mg/mL and spread on an indium tin oxide (ITO)-coated glass slide (50 mm ×50 mm, 10-15 Ω/in^2^; Adafruit, Brooklyn, NY, P1310) before desiccating the slides under vacuum for 1 hour. Copper tape (Adafruit P1128) was adhered to the conductive side of the slides, and a closed chamber was formed using a Teflon spacer and binder clips. The chamber was then filled with 100 mM sucrose. The copper tape electrodes were connected to a voltage generator and a sine wave current of 0.1 Vpp, 10 Hz, was applied, increasing the voltage by 0.1 Vpp every two minutes until 1.5 Vpp was reached. The current was continued for 2 hours at 1.5 Vpp before GUVs were harvested. GUVs were used within 24 hours of formation and were stored at 4°C. All GUV preparations were done without direct exposure to light to prevent oxidation of phospholipids.

### Zeta potential testing

SUV zeta potential was measured on a Malvern Zetasizer Nano-ZS, utilizing a Zetasizer universal dip cell (ZEN1002) at 25°C.

### GUV imaging

GUVs formed in sucrose solution were diluted in hypertonic glucose solution. The glucose concentration was 4 mOsm/L higher than the sucrose, with osmolarity verified on an Advanced Instruments osmometer, model 3320. Vesicles were imaged on a Nikon Ti-E inverted microscope equipped with a Ph3 phase ring, CFI Plan Apochromat 100x oil immersion objective (numerical aperture 1.45, working distance 0.13 mm), Nikon DS-Qi2 camera and NIS Elements v.4.20.01 for image acquisition. Images were taken every 300 ms for 5 minutes, with an exposure time of 30 ms per frame (see Supplemental Movies).

### GUV edge extraction algorithm

Edge detection algorithms that identify vesicle contours using local extrema in image intensity or intensity gradients (e.g. Refs. (21–25)) often fail when vesicle images contain contaminating structures such as neighboring vesicles, imaging artifacts, or localized bright/dark regions. Consequently, images containing such defects are frequently discarded prior to analysis. Because the majority of images in our dataset contained these features, we developed a geometry-informed edge extraction algorithm that iteratively constrains the feasible contour-search region in polar coordinates. The algorithm exploits the expectation that vesicle contours are smooth, approximately circular, and dominated by low-order angular fluctuations, allowing reliable edge identification even in the presence of substantial image defects.

We have provided a detailed edge extraction protocol in the *Supplemental Methods* section for reference, but provide a brief overview here. Each image is represented as a two-dimensional array of pixel intensities, *I*_*i, j*_ (Fig. 2A). Our algorithm uses the following four-step process for each vesicle image: (1) identification of the approximate vesicle center, (2) mapping the image to polar coordinates, (3) iterative reduction of the feasible contour-search region, and (4) post-process filtering.

**Figure 2:**
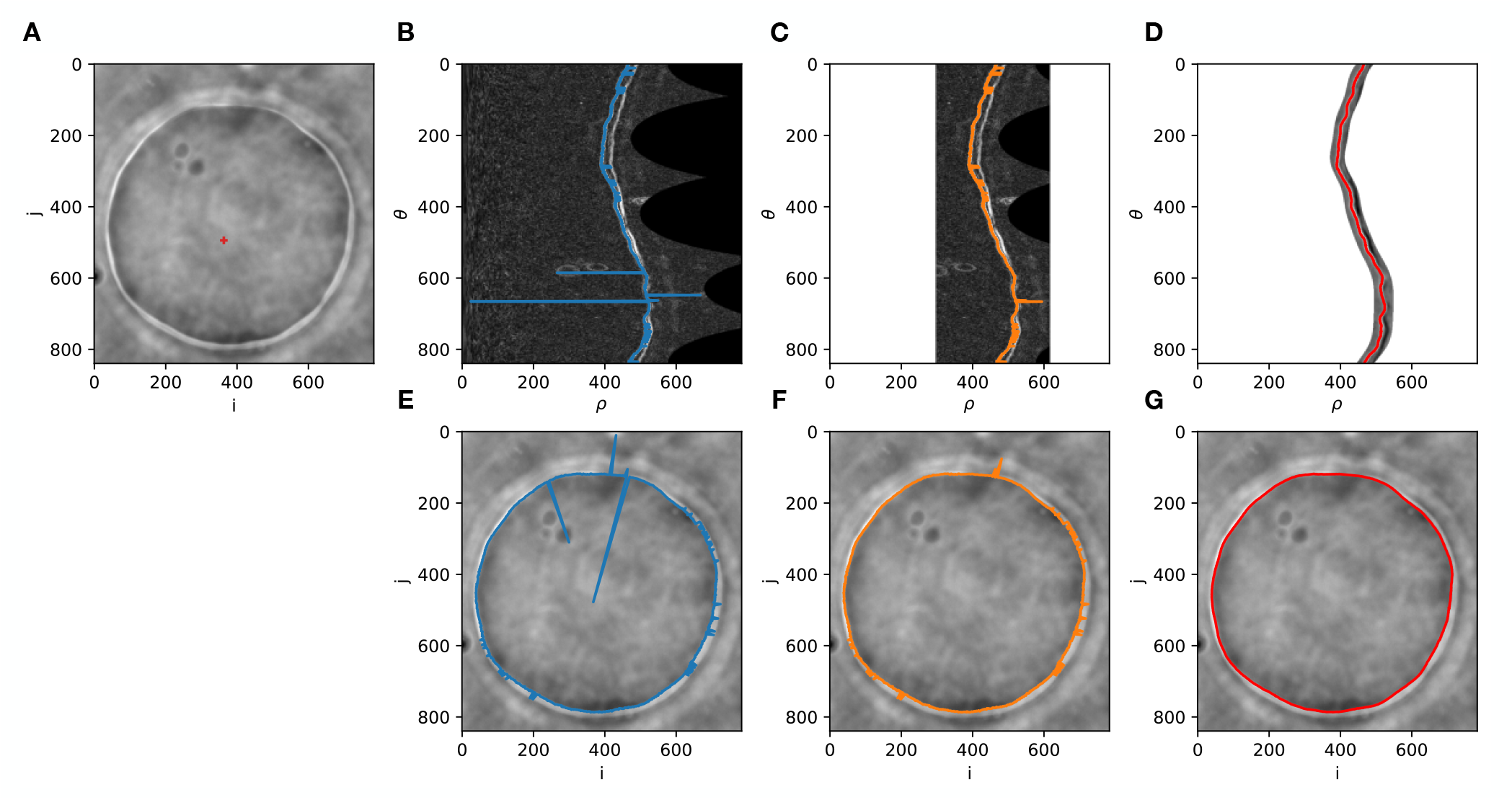
GUV edge extraction steps. *A)* Intensities *I*_*i, j*_ from a single frame of a vesicle video taken via phase contrast microscopy. The approximate vesicle center (*x*_0_, *y*_0_) is shown with a red mark. *B) S*(*ρ, θ*), the image after mapping to polar coordinates and applying a Sobel filter. *ρ*\*(*θ*) (blue line) is the pixel of greatest gradient magnitude in each angular bin. *C)S*\*(*ρ, θ*) is the same image as in *B*, but with the preliminary mask applied. *ρ*\*\*(*θ*) (orange line) is the pixel of greatest gradient magnitude in each angular bin after masking. *D) I*\*(*ρ, θ*) is the original image after mapping to polar coordinates and applying the Fourier-informed mask. The detected vesicle edge *ρ*(*θ*) (red line) is overlaid. *E-G)* Original image *I*_*i, j*_ is shown with *ρ*\*(*θ*)/*C* (panel *E*); *ρ*\*\*(*θ*)/*C* (panel *F*); and *r* (*θ*) (panel *G*) overlaid, respectively.

In step 1, the vesicle’s center coordinate (*x*_0_, *y*_0_) is approximated by applying a Sobel filter (26) to highlight sharp gradients in pixel intensity, blurring the filtered image, separating foreground and background pixels via Otsu’s method (27), and then measuring the center of mass of the foreground pixels. Step 2 maps the pixels in *I*_*i, j*_ to polar coordinates *I* (*ρ, θ*) around (*x*_0_, *y*_0_), where *ρ* ≡ *r*/*C* is the radial distance *r* from (*x*_0_, *y*_0_) scaled by *C*, a term determined by the area of *I*_*i, j*_ included during the transformation to *I* (*ρ, θ*). Here, we included all pixels of *I*_*i, j*_ within half the image diagonal of (*x*_0_, *y*_0_), so 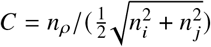, where *n*_*i*_, *n* _*j*_, and *n*_*ρ*_ are the maximal indices in the *i, j*, and *ρ* dimensions of their respective images.

Step 3 involves iteratively masking regions of *I* (*ρ, θ*) that do not correspond to the vesicle edge before extracting the best edge candidate for further refinement. A Sobel filter is applied to *I* (*ρ, θ*) to yield *S*(*ρ, θ*) in order to highlight sharp gradients in pixel intensity. We first calculate *ρ*\* (*θ*) by finding the *ρ* position corresponding to the pixel of maximum intensity in each *θ* row of *S* (*ρ, θ*). This quantity is comparable to what most existing edge extraction algorithms return, but as shown in Fig. 2B and 2E, it does not reliably follow the true vesicle edge when visual anomalies are present. We then mask all *ρ* outside of some tolerance from ⟨*ρ*\* (*θ*) ⟩ _*θ*_ and again find the pixel of maximum intensity in each row, *ρ*\*\* (*θ*). As shown in Fig. 2C and 2F, this preliminary mask is insufficient to isolate the vesicle edge in all cases. We next Fourier transform *ρ*\*\* (*θ*), suppress all but the lowest modes, and inverse Fourier transform to yield a denoised signal *ρ*^†^(*θ*) from which to construct a new mask and apply it to *I* (*ρ, θ*) (Fig. 2D). This now-masked *I*\* (*ρ, θ*) has a Sobel filter and Gaussian filter applied along the *ρ* dimension, before the pixel of maximum intensity is chosen in each row to represent the detected edge *ρ* (*θ*) (red line, Fig. 2D). Dividing *ρ* (*θ*) by *C* yields the true vesicle edge *r* (*θ*), devoid of any radial distortion incurred during the transformation to polar coordinates in step 2 (Fig. 2G).

In the rare event that nearby vesicles or excessive imaging artifacts drastically bias the estimation of (*x*_0_, *y*_0_), steps 1-3 may fail to isolate a complete edge. In these cases, detected edges will often feature sharp discontinuities in *r* (*θ*) that can be filtered by measuring the curvature in *r* with respect to *θ*. In step 4 we search for failures in the process by rejecting any detected edge that has curvature in excess of a threshold value.

The edge extractor was implemented in Python 3.11 using the external libraries *scipy*.*ndimage, skimage*.*filters, skimage*.*measure, cv2*, and *numpy*. The images in this study were read into numpy arrays using the *nd2* library. Our codebase is available in a public GitHub repository (28).

### Analysis of GUV fluctuations

Vesicle edges detected using the above algorithm *r* (*θ, t*) are interpolated onto a common angular grid with 120 bins (Δ*θ* = π /60) to ensure consistent angular sampling across vesicles, yielding *R* (*θ, t*). The average radius of each vesicle, *R*_0_ ≡ ⟨*R*(*θ, t*) ⟩ _*θ,t*_, was used to normalize radial fluctuations 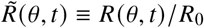. The Discrete Fourier Transform of 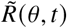 is defined as

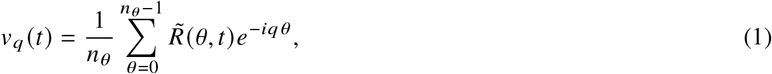

where *v*_*q*_ (*t*) are the amplitudes for each integer Fourier mode *q*, and *n*_*θ*_ = 120 angular bins. The amplitudes are multiplied by their complex conjugate and averaged over time to yield ⟨|*v*_*q*_ |^2^ ⟩_*t*_. Positive Fourier modes 3 through 8 are then fit using the non-linear least-squares method to the functional form adapted from Ref. (22),

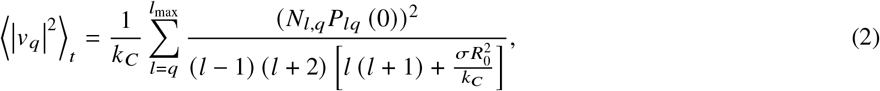

where *k*_*C*_ is the bending modulus, *Nl,q* is the normalization constant 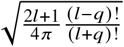, *Plq* (0) is the associated Legendre polynomial evaluated at 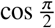, and *σ* is the surface tension. In this fitting procedure *k*_*C*_ and *σ* are free fit parameters, and *l*_max_ is set to 500. The spectrum over the range *q* = 3 to *q* = 8 was observed to be insensitive to further increases in *l*_max._

Vesicles with excess surface tension (|*σ*| ≥*σ*_max_) are typically removed from analysis (e.g. see Refs. (25, 29)) in order to ensure that surface tension does not dominate too much of the power spectrum, with 10^−8^ N/m representing one such recommended *σ*_max_ value (29). To determine which *σ*_max_ threshold to use when extracting *k*_*C*_, we measured *k*_*C*_ for 100% DOPC vesicles while altering *σ*_max_ (see Fig. S2). When *σ*_max_ = 10^−8^ N/m is used, the measured *k*_*C*_ is 26 ± 7 *k* _*B*_*T* (*n* = 8), equaling the *k*_*C*_ obtained in Ref. (14). As *σ*_max_ is increased, the measured *k*_*C*_ increases monotonically, reaching 74 *k* _*B*_*T* (*n* = 31) when *σ*_max_ = 10^−5^ N/m. We were unable to isolate enough sphingolipid-containing vesicles at the lower *σ*_max_ thresholds to perform our analysis (see Fig. 3), so we chose *σ*_max_ = 1.325 ×10^−7^ N/m because it allowed for collection of at least 5 replicas per data point while minimizing the measured *k*_*C*_ for pure DOPC to within error of 26 *k* _*B*_*T*. This *σ* threshold corresponds to an allowable vesicle surface area expansion of less than 5 × 10^−5^ percent, where percent surface area expansion is defined as *σ*/*k* _*A*_ × 100 and the measured area compressibility modulus *k* _*A*_ of DOPC is 0.265 N/m (12).

**Figure 3:**
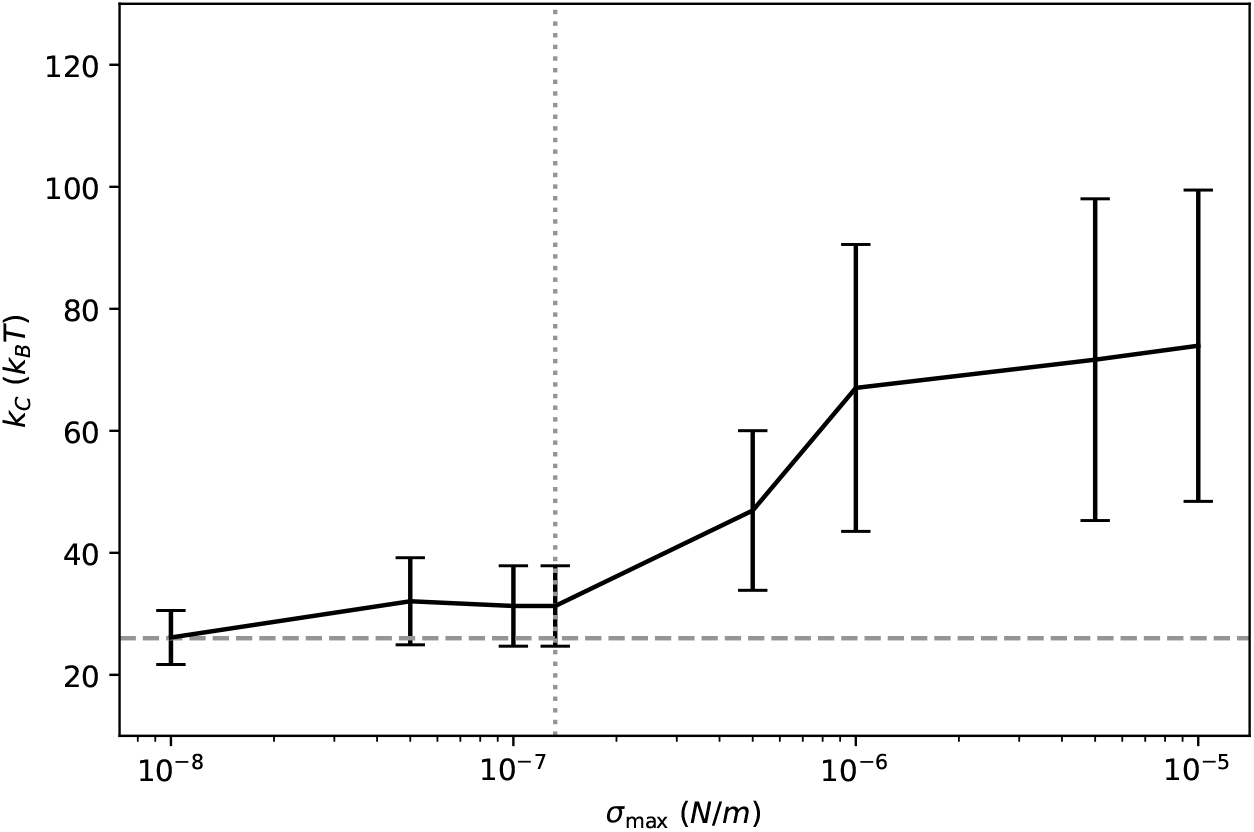
Bending modulus *k*_*C*_ for DOPC vesicles as surface tension threshold *σ*_max_ varies. Dotted vertical line denotes threshold used for all measurements in Fig. 5. All errors are reported as standard error of the mean with *n* ≥ 8 replicas. The reference value for 100% DOPC (14) is shown as a horizontal dashed gray line.

After filtering out vesicles with excess surface tension, the amplitudes of each mode are then averaged together for all replicas of a particular vesicle composition. The positive Fourier modes 3 through 8 of this average spectrum are fit to the functional form above with *k*_*C*_ as the sole fit parameter and *σ* = 0. Standard error of the mean is reported for each replica series, with *n* ≥ 5. Fluctuation analysis code is available in a public GitHub repository (28), and utilizes Python 3.11 with numpy (30) and lmfit (31).

## RESULTS AND DISCUSSION

To understand the impact of CPG on biological and artificial membranes, we first purified CPG from *C. crescentus* membrane extracts (see Materials and Methods). The purified CPG was then incorporated into phospholipid liposomes for characterization.

### CPG increases the magnitude of liposomal zeta potential

The stability of membrane vesicles in solution is partially dependent on their potential for aggregation. To assess the effects of CPG on liposome aggregation, we measured the zeta potential, which reflects the charge that is carried with the membrane. This is different from surface charge as zeta potential accounts for the ions in solution that interact with the membrane, while surface charge is a measure of solely the charge of the particles within the membrane. As like charges repel each other, zeta potential is an indicator of the stability of particles within solution; it does not determine the stability of individual particles, but formulations with higher zeta potential are less likely to aggregate. Zeta potential was determined on POPC-based small unilamellar vesicles (SUVs) containing varying amounts of neutral and anionic sphingolipids.

Of the three sphingolipids tested, C1P had the greatest magnitude of zeta potential when measured in PBS containing Mg^2+^ and Ca^2+^ ions (Fig 4A). By contrast, the zeta potential of vesicles containing ceramide or CPG were similar and of lower magnitude than C1P. We hypothesized that the lower zeta potential of CPG may be due to the phosphate in PBS interacting with the phosphate in CPG, but when these experiments were repeated in HEPES buffer we obtained similar results. These data demonstrated that the inclusion of any of the sphingolipids tested increased the charge of the particles; C1P liposomes have a greater magnitude of zeta potential and are thus more likely to remain stable without aggregation compared to the other two sphingolipids tested.

**Figure 4:**
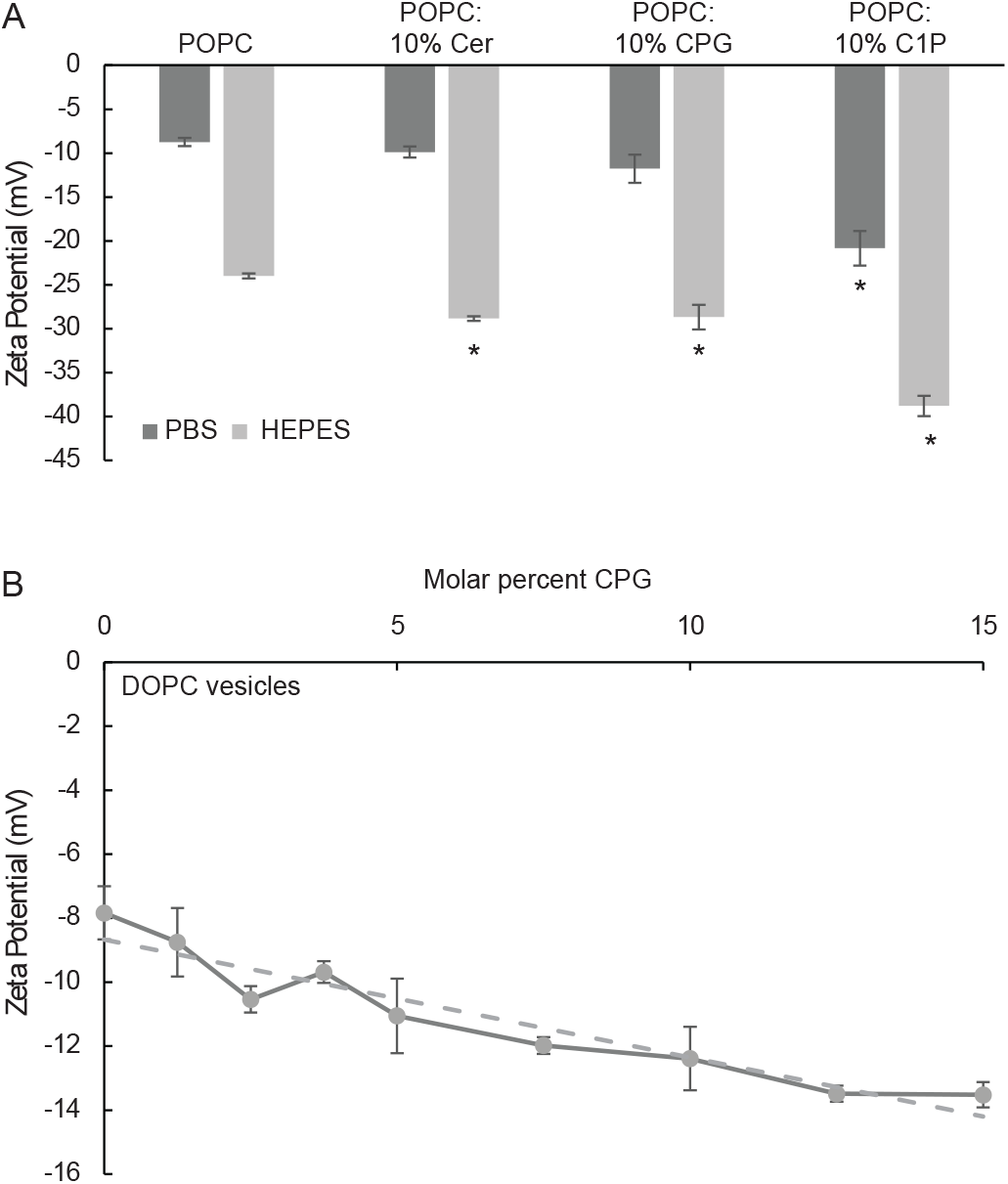
Zeta potential of SUVs containing CPG. (A) SUVs were made from POPC and either ceramide, CPG, or C1P, 90:10 w/w. SUVs made in PBS with Mg^2+^/Ca^2+^ had a consistently lower zeta potential than those made in HEPES buffer, indicating that HEPES buffer is less likely to cause aggregation. While the inclusion of any ceramide increased the magnitude of the zeta potential, C1P had the greatest impact in both PBS and HEPES. (n = 3; error bars are SE, ANOVA_PBS_ F (3, 11) = 17.13, p = 7.65 ×10^−4^; ANOVA_HEPES_ F (3, 11) = 45.03, p < 0.0001; * post hoc comparisons to POPC using Tukey test, p < 0.05). (B) SUVs containing DOPC and increasing molar fractions of CPG were then analyzed for zeta potential, and a linear equation for the slope indicated a decrease of 0.37 mV per mole percent CPG (*R*^2^ = 0.91).

We further investigated the impact of CPG on SUVs based on DOPC. When increasing concentrations of CPG were incorporated into the DOPC liposomes, we observed a linear decrease of the zeta potential (Fig 4B).

### CPG increases membrane bending stiffness

To understand how the addition of neutral ceramide to a homogeneous phospholipid membrane changes the flexibility of the binary mixture, we determined the bending modulus *k*_*C*_ via flicker spectroscopy of vesicles containing varying amounts of ceramide and DOPC (Fig. 5). As described in Methods, we included vesicles with higher surface tension (*σ*≤ 1.325 × 10^−7^ N/m) in our analysis to facilitate collection of at least 5 replicas per lipid concentration. The DOPC membrane without ceramide has a *k*_*C*_ of 31 ±7 *k* _*B*_*T*, which is within error of established *k*_*C*_ measurements of pure DOPC (14), although we are able to obtain a measurement of 26 ±7 *k* _*B*_*T* when we remove vesicles with *σ* > 10^−8^ N/m. Increasing the amount of ceramide resulted in a non-monotonic transition in *k*_*C*_, ranging from 58± 6 *k* _*B*_*T* at 2.5% ceramide to 19 ±5 *k* _*B*_*T* at 7.5% ceramide. There was no significant difference in *k*_*C*_ between a vesicle with 10% ceramide, 5% ceramide, and a pure DOPC vesicle. The wide variance in *k*_*C*_ at different amounts of ceramide could be explained by the complex phase behavior of neutral ceramides as they are known to form domains in POPC at similar temperatures and concentrations to those used here (32, 33), suggesting that the complex results may be due to membrane reorganization with increasing ceramide concentrations.

**Figure 5:**
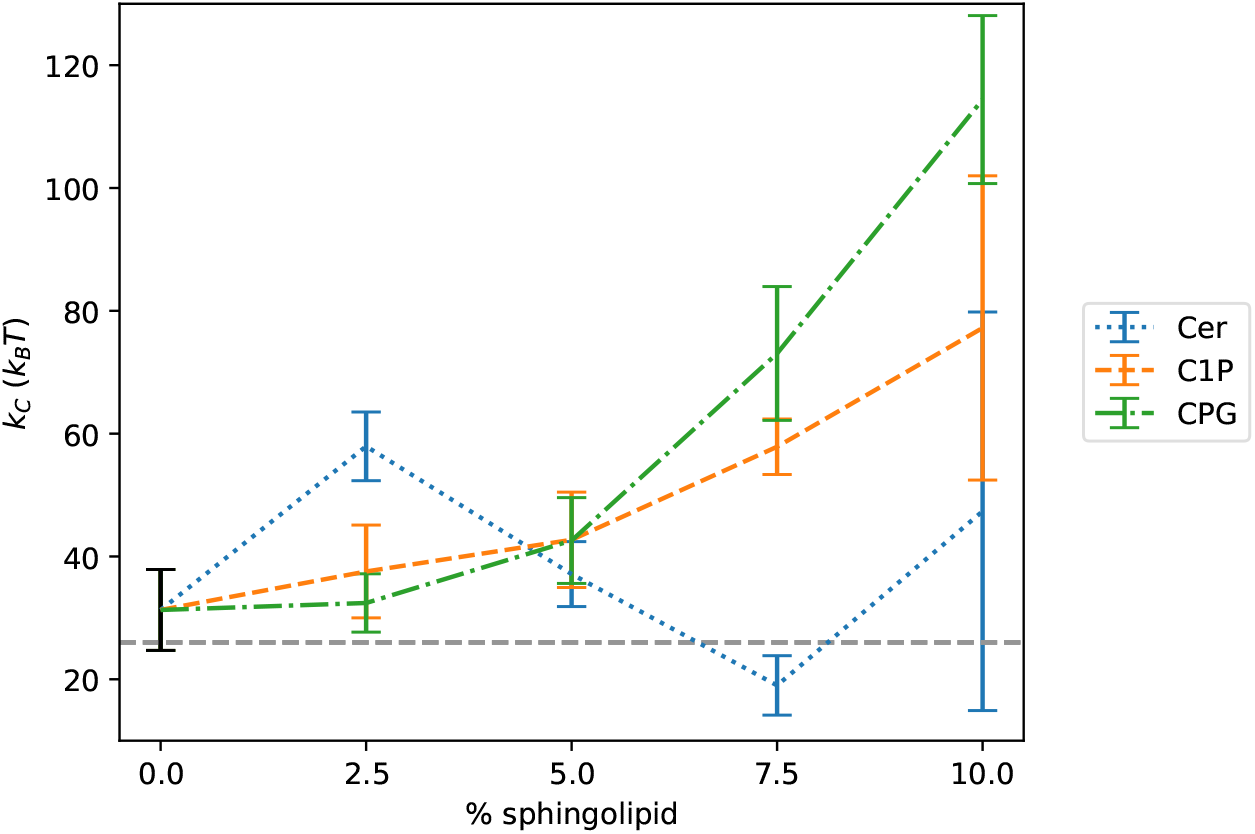
Bending modulus *k*_*C*_ for DOPC vesicles containing varying amounts of sphingolipid. Bending modulus of pure DOPC vesicles appears in black. The reference value for 100% DOPC (14) is shown as a horizontal dashed gray line. Error is represented as standard error of the mean with *n* ≥5 replicas. Vesicles with| *σ*| 1.325 ×10^−7^ N/m are removed from the analysis (see Methods).

To assess whether the addition of sphingolipids containing a charged headgroup changes the membrane flexibility, we determined the bending modulus of DOPC vesicles with varying amounts of the anionic sphingolipids C1P and bacterially-derived CPG (Fig. 5). *k*_*C*_ increases with the percentage of either C1P or CPG in the vesicle, reaching values in excess of 60 *k* _*B*_*T* at 10% anionic sphingolipid. At mole fractions exceeding 5% sphingolipid, vesicles containing CPG are significantly stiffer than those containing C1P. This suggests that the presence of glycerate in the head and/or a hydroxyl group in the tail of the bacterially-derived CPG may enhance the stiffening effect of anionic sphingolipids. The membrane stiffening observed upon addition of anionic lipid is consistent with, but greater in magnitude than, existing findings (25) that show charged headgroups stiffen membranes. When compared to the non-monotonic behavior observed when adding neutral ceramide, our findings suggest that the effect of headgroup charge dominates even despite significant differences in tail and backbone chemistry between DOPC and sphingolipids. Our findings also indicate (see Fig. S2) that varying the surface tension threshold *σ*_max_ does not shift the relative differences in *k*_*C*_ across the ceramide species and concentrations, with the possible exception of 10% C1P. This suggests that the decision to use a less-stringent cutoff (see Methods) may introduce a slight systematic error of a few *k* _*B*_*T*, but that the trends we observe are robust to the choice of cutoff.

## CONCLUSION

This work provides methodological advances that may facilitate future investigations of membrane mechanics in bacterial and charged lipid systems. We developed a purification protocol for charged bacterial lipids that yields material suitable for quantitative biophysical characterization, and we introduced an edge-detection workflow designed to mitigate common imaging artifacts that can otherwise reduce the number of analyzable vesicles. Together, these advances expand the range of experimental systems that can be brought to bending rigidity measurements. To support reproducibility and broader adoption, all protocols and analysis software have been made publicly available.

Having determined that the inclusion of ceramides increases the membrane stiffness in liposomes, it leads us to question whether this impact would be observed *in vivo*. In *Escherichia coli*, the outer membrane is a significant contributor to the overall mechanical stiffness of the cell (34). Considering that sphingolipids, including CPG, are found in the outer membrane (17) and that CPG increases the bending stiffness of liposomes, we hypothesize that the production of CPG by *C. crescentus* may contribute to the rigidity of the outer membrane and therefore the strength of the cell. The production of these anionic sphingolipids could help maintain cellular integrity in the hypo-osmotic conditions commonly found in *C. crescentus*’s freshwater ecological niche.

Beyond microbial physiology, when considering the broader impacts of sphingolipids on vesicle membranes, we also considered whether these novel lipids would be useful in lipid nanoparticle (LNP) drug delivery systems. LNPs are commonly used to administer therapeutics that are unstable (*e*.*g*., RNA vaccines) or harmful to the body in higher doses (*e*.*g*., chemotherapeutics). Sphingolipids have been shown to promote longer *in vivo* stability and increased blood circulation in LNPs, though no LNP drugs currently utilize sphingolipids as membrane components (35). CPG has been demonstrated to decrease zeta potential, which indicates that liposomes containing CPG would be less likely to aggregate after formation. The increased membrane stiffness indicates that these liposomes would be less likely to leak their contents. More broadly, the discovery of novel bacterial lipids with uncharacterized chemical and biophysics properties may be a useful source of novel membrane components for future LNP development projects.

## AUTHOR CONTRIBUTIONS

J. Chamberlain purified CPG and performed all experimental methods. J. Sandberg and B. Bratton designed the edge extractor algorithm. J. Sandberg conducted the mathematical analysis to calculate liposomal bending stiffness. Z. Guan performed all LC/MS analysis. G. Brannigan and E. Klein consulted on experimental design and execution. J. Chamberlain and J. Sandberg wrote the original manuscript. E. Klein and G. Brannigan edited the manuscript.

## ACKNOWLEDGMENTS

We thank Julianne Griepenburg and Jesse Cecero (Rutgers University-Camden) for assistance in generating SUVs. We thank Tanisha Dhakephalkar (Rutgers University-Camden) for assistance in cloning. We thank Petia Vlahovska and Diptendu Sen (Northwestern University) for assistance in performing flicker spectroscopy.

This work was funded by NSF grants MCB-2031948 (to EAK) and MCB-2224195 (to EAK and GB).

