## Supplement for "Anionic bacterial sphingolipids increase membrane stiffness"

**Supplemental Materials**

### 1 Supplemental methods

Each image is represented as a two-dimensional array of pixel intensities, $I_{i,j}$ (Fig. 2A). Our algorithm then uses the following four-step process for each vesicle image: (1) identification of the approximate vesicle center, (2) mapping the image to polar coordinates, (3) iterative reduction of the feasible contour-search region, and (4) post-process filtering.

1. Identify the approximate center of the vesicle.
   (a) Highlight sharp intensity gradients between neighboring pixels by applying a Sobel filter [2] to $I_{i,j}$ and measuring its magnitude to obtain $S_{i,j}$.
   (b) Reduce background noise by applying a Gaussian filter $G_{i,j} = \mathcal{G}_\sigma[S_{i,j}]$, where $\mathcal{G}_\sigma$ is a Gaussian kernel with standard deviation $\sigma$. Here we used $\sigma = 10$.
   (c) Using Otsu's method [1], calculate the threshold intensity $G^*$ that best distinguishes foreground from background pixels of $G_{i,j}$.
   (d) Find the set $H$ of all pixels satisfying $G_{i,j} > G^*$.
   (e) Estimate the coordinates of the vesicle center using $x_0 = \langle H \rangle_x$, and $y_0 = \langle H \rangle_y$.

2. Resample the Cartesian image $I_{i,j}$ onto a discretized polar grid,

   $$I_{\alpha,\beta} = I(x_0 + \rho_\alpha \cos\theta_\beta,\, y_0 + \rho_\alpha \sin\theta_\beta),$$

   where $\rho_\alpha = C\alpha\Delta\rho$ and $\theta_\beta = \beta\Delta\theta$ denote discretized radial and angular coordinates centered on $(x_0, y_0)$. $\Delta\rho$ and $\Delta\theta$ are the bin widths in $\rho$ and $\theta$, respectively, and $C$ is the constant scaling factor relating the transformed radial coordinate $\rho$ to the true radial distance $r$.

   Here, we used *cv2.linearPolar*, which resamples the Cartesian image onto a polar grid using bilinear interpolation. The output image was generated with the same dimensions as $I_{i,j}$, consequently

   $$C = \frac{n_\rho}{\frac{1}{2}\sqrt{n_i^2 + n_j^2}},$$

   where the denominator is the Euclidean distance from the image center to the image corner, $n_\rho$ is the maximum sampled radial coordinate, and $n_i$ and $n_j$ are the maximum Cartesian image indices. Coordinates outside the bounds of the original Cartesian image were assigned intensity 0.

   For clarity, subsequent operations on the transformed polar image are written as functions of the sampled coordinates $\rho$ and $\theta$ rather than the explicit indices $\alpha$ and $\beta$.

3. Iteratively reduce the feasible contour-search region.
   (a) Preliminary contour estimate.
      i. Compute $S(\rho, \theta)$, the Sobel gradient magnitude of $I(\rho, \theta)$.
      ii. Measure $\rho^*(\theta)$, the radial position of the maximum-gradient pixel in each angular bin of $S(\rho, \theta)$: $\rho^*(\theta) \equiv \underset{\rho}{\operatorname{argmax}}\, S(\rho, \theta)$.
      This quantity is comparable to what most existing edge extraction algorithms return, but as shown in Fig. 2B and 2E, it does not reliably follow the true vesicle edge when visual anomalies are present.
      iii. Compute the mean radial position of the maximum-gradient pixels, $\langle \rho^*(\theta) \rangle_\theta$.

iv. Because vesicle contours are expected to deviate only modestly from circularity, regions far inside or outside the mean detected radius can be excluded without affecting the detected contour. Construct the mask $S^*(\rho, \theta)$:

$$S^*(\rho,\theta) = \begin{cases} 0, & \rho \leq \frac{3}{4}\langle \rho^*(\theta)\rangle_\theta \\ 0, & \rho \geq \frac{5}{4}\langle \rho^*(\theta)\rangle_\theta \\ S(\rho,\theta), & \text{otherwise.} \end{cases}$$

v. Measure $\rho^{**}(\theta)$, the radial position of the maximum-gradient pixel in each angular bin of $S^*(\rho, \theta)$: $\rho^{**}(\theta) \equiv \underset{\rho}{\text{argmax}}\, S^*(\rho, \theta)$.

As shown in Fig. 2C and 2F, the preliminary mask and corresponding contour estimate $\rho^{**}(\theta)$ remain susceptible to spurious edge detections arising from imaging artifacts and/or neighboring structures. We therefore exploit the expectation that vesicle contours are smooth and dominated by low-order angular fluctuations to further constrain the feasible contour-search region.

(b) Fourier-informed contour refinement.

i. Compute the Fourier amplitudes $A(q')$ of $\rho^{**}(\theta)$ using a one-dimensional Discrete Fourier Transform (DFT):

$$A(q') = \sum_{\theta=0}^{n_\theta - 1} \rho^{**}(\theta) e^{-iq'\theta}.$$

ii. High-order angular fluctuations are assumed to arise predominantly from spurious edge detections and imaging artifacts at the spatial scales resolved in these images; consequently, only low-order Fourier modes were retained. Mask the amplitudes $A(q')$ as follows:

$$A^*(q') = \begin{cases} 0 + 0i, & q' < 0 \\ 0 + 0i, & q' \geq 7 \\ A(q'), & \text{otherwise.} \end{cases}$$

iii. Take the inverse DFT of $A^*(q')$ to obtain $\rho^\dagger(\theta)$:

$$\rho^\dagger(\theta) = \frac{1}{n_\theta} \sum_{q'=0}^{n_\theta - 1} A^*(q') e^{iq'\theta}.$$

High-frequency spurious edge detections are substantially suppressed in $\rho^\dagger(\theta)$, yielding a smooth approximation of the vesicle contour.

iv. Construct the masked image $I^*(\rho, \theta)$ by applying the following mask independently within each angular bin:

$$I^*(\rho,\theta) = \begin{cases} 0, & \rho \leq \frac{19}{20}\rho^\dagger(\theta) \\ 0, & \rho \geq \frac{21}{20}\rho^\dagger(\theta) \\ I(\rho,\theta), & \text{otherwise.} \end{cases}$$

The resulting image retains only a narrow radial band spanning $\pm 5\%$ around the smooth contour estimate $\rho^\dagger(\theta)$, as shown in Fig. 2D.

(c) Local contour refinement.

i. Apply a one-dimensional Sobel operator $\mathcal{S}$ along the $\rho$ dimension of $I^*(\rho, \theta)$ to yield $S_\rho(\rho, \theta)$: $S_\rho(\rho, \theta) = \mathcal{S}_\rho[I^*(\rho, \theta)]$.

ii. Apply a one-dimensional Gaussian kernel $\mathcal{G}_\sigma$, where $\sigma$ is the standard deviation, along the $\rho$ dimension of $S_\rho(\rho, \theta)$ to yield $G_\rho(\rho, \theta)$: $G_\rho(\rho, \theta) = \mathcal{G}_{\rho,\sigma}[S_\rho(\rho, \theta)]$. Here we used $\sigma = 2$.

iii. Measure the detected contour $\rho(\theta) \equiv \underset{\rho}{\operatorname{argmax}}\, G_\rho(\rho, \theta)$.

(d) Convert the detected contour to the true radius using $r(\theta) = \rho(\theta)/C$, eliminating any radial distortion created during the transformation to polar coordinates.

4. Filter out bad edge detections by detecting high curvature.

(a) Measure the curvature of the detected edge $\Delta^2_\theta r(\theta)$. Here we used a finite second difference where $\Delta^2_\beta r(\theta_\beta) = r(\theta_{\beta+1}) - 2r(\theta_\beta) + r(\theta_{\beta-1})$.

(b) Filter out vesicles with high curvature. Here, $\max|\Delta^2_\theta r(\theta)| > 5$ were excluded empirically.

### 2 Supplemental Figures

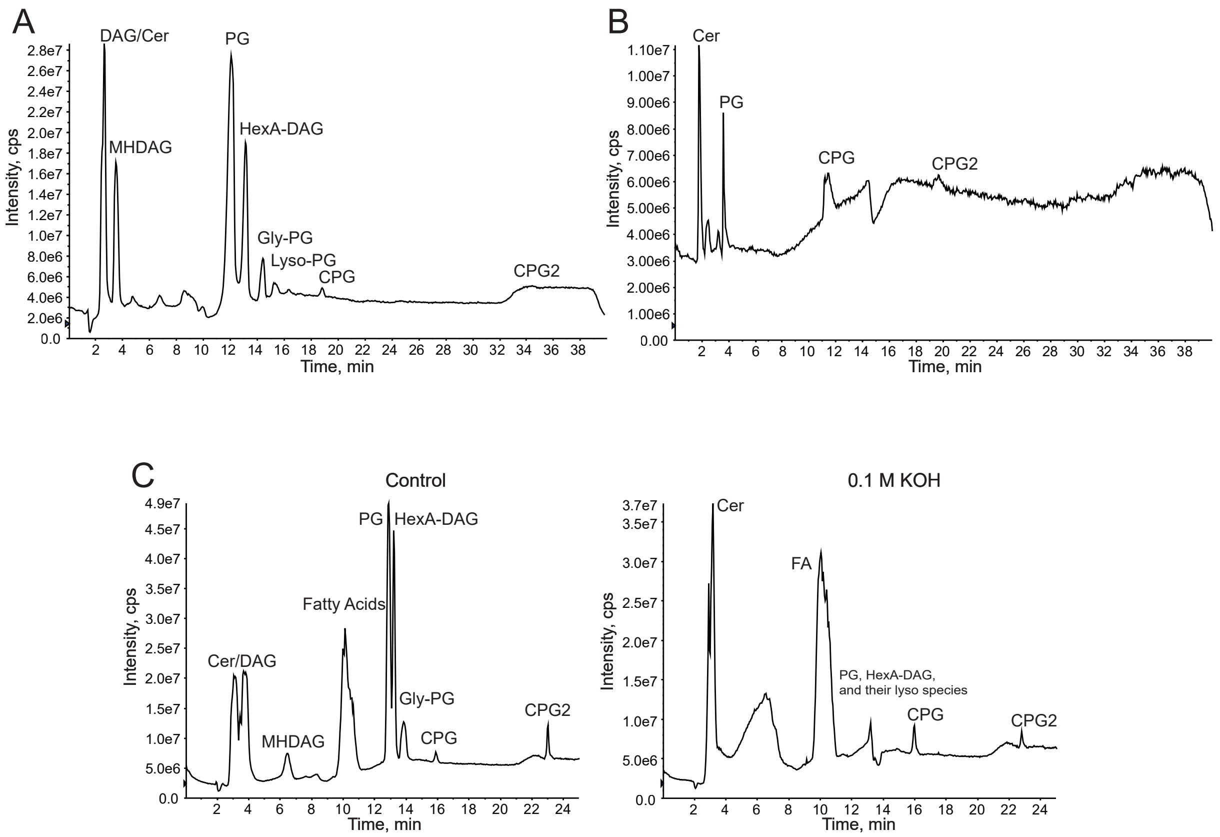


Figure S1: Lipid extraction techniques to enrich and isolate novel bacterial sphingolipids. (A) LC/MS data of a typical lipid extraction by the method of Bligh and Dyer, where only negligible amounts of CPG and CPG2 are purified. (B) LC/MS and MS/MS analysis of an acidified lipid extraction. By acidifying the aqueous phase before isolating the lower phase, it is possible to enrich the amounts of CPG that are purified. (C) LC/MS analysis of lipid extracts after exposing them to 0.1 M KOH for 4 hours demonstrates that mild alkaline hydrolysis degrades the phospholipids into free fatty acids. These fatty acids are more easily removed by preparative TLC. The TLC is run in a $CHCl_3$:MeOH:acetic acid:$H_2O$ 8:3:2:1 solvent system, and the lipids can be scraped off to yield a highly enriched sample of CPG (See Figure 1B).

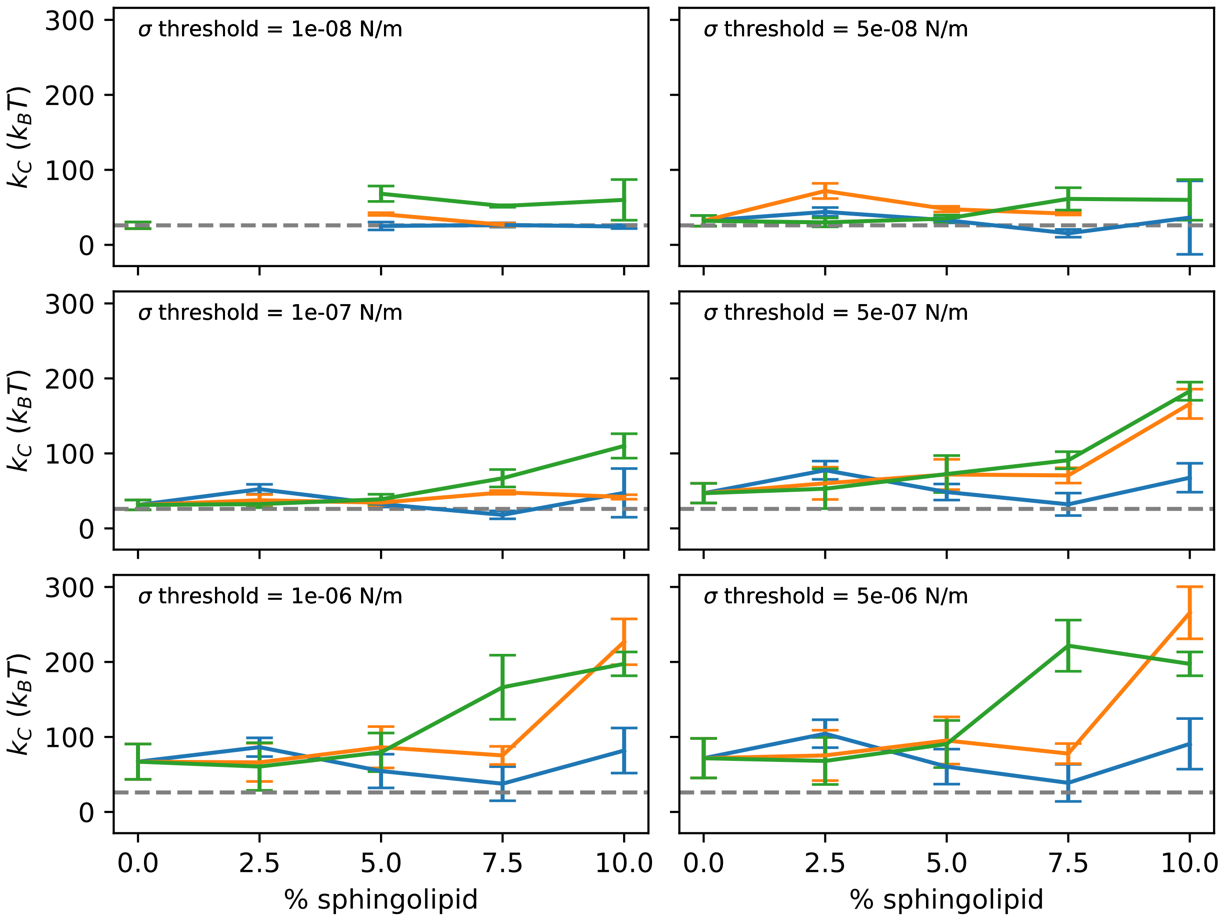


Figure S2: Bending modulus $k_C$ for DOPC vesicles containing varying amounts of sphingolipid, measured at different surface tension thresholds $\sigma_{\max}$. Data points for ceramide (orange line), C1P (blue line), and CPG (green line) are shown when there were more than two samples obtained.

#### 3 Supplemental videos

Movies S1-S4 are examples of time-lapse videos of fluctuating vesicles. S1 DOPC, S2 DOPC with 10% ceramide, S3 DOPC with 10% C1P, and S4 DOPC with 10% CPG.

Movies S5-S8 are the edge detection videos corresponding to the vesicles shown in Movies S1-S4.

[2] Irwin Sobel. An isotropic 3x3 image gradient operator. *Presentation at Stanford A.I. Project 1968*, 1968.
